# Loop 1 of human cardiac myosin regulates the rate of ATP induced actomyosin dissociation but not the rate of ADP release

**DOI:** 10.1101/2021.10.08.463735

**Authors:** Akhil Gargey, Yuri E. Nesmelov

## Abstract

Double mutation D208Q:K450L was introduced in the beta isoform of human cardiac myosin to remove the salt bridge D208:K450 connecting loop 1 and the seven-stranded beta sheet within the myosin head. Beta isoform-specific salt bridge D208:K450 was previously discovered in the molecular dynamics simulations. It was proposed that loop 1 modulates nucleotide affinity to actomyosin and we hypothesized that the electrostatic interactions between loop 1 and myosin head backbone regulates ATP binding to and ADP dissociation from actomyosin, and therefore, the time of the strong actomyosin binding. Wild type and the mutant of the myosin head construct (1–843 amino acid residues) were expressed in differentiated C2C12 cells, and the kinetics of ATP induced actomyosin dissociation and ADP release were characterized using transient kinetics spectrophotometry. Both constructs exhibit a fast rate of ATP binding to actomyosin and a slow rate of ADP dissociation, showing that ADP release limits the time of the strongly bound state of actomyosin. We observed a faster rate of ATP-induced actomyosin dissociation with the mutant, compared to the wild type actomyosin. The rate of ADP release from actomyosin remains the same for the mutant and the wild type actomyosin. We conclude that the flexibility of loop 1 is a factor affecting the rate of ATP binding to actomyosin and actomyosin dissociation. We observed no effect of loop 1 flexibility on the rate of ADP release from actomyosin.

**Highlights:**

1. Human cardiac myosin has the isoform-specific salt bridge, electrostatically linking loop 1 and myosin head backbone.
2. Absence of loop 1 – backbone salt bridge increases rate of ATP-induced actomyosin dissociation
3. The rate of ADP dissociation from actomyosin is not affected by the isoform-specific salt bridge.

## Introduction

Myosins are a family of molecular motors, transforming the chemical energy of ATP into mechanical motion of muscles and cells. Myosin head interacts with actin and nucleotide and changes conformation two times in a cycle. The cyclic conformational change of the head results in the production of movement and force. Myosins have different functions in cells and organisms, and therefore, have different kinetic properties. The kinetic differences are determined by the amino acid sequence, which governs protein structure and interactions between structural elements. The sequence analysis suggests that two surface loops of the myosin head, loop 1 and loop 2, likely have regulatory functions, since the sequence variability of these loops significantly exceeds the sequence variability of the core myosin (1). Loop 2 is located within the actin binding interface and the chimera studies confirmed that loop 2 regulates myosin interaction with actin (2). Loop 1 is located near the active site, and hypothetically regulates interaction with nucleotide, most likely ADP release (3), and therefore, regulates the velocity of muscle shortening. Loop 1 links two helices of the myosin head, one connecting to loop switch 1 and another to P-loop of the nucleotide binding site (Figure 1). P-loop, or Walker A motif, coordinates Mg ion, complexed with ATP within the active site, and stabilizes ATP during hydrolysis. Loop switch 1 coordinates Mg ion and γ phosphate of ATP. This proximity to the loops at the active site makes loop 1 a hypothetical regulatory element, controlling nucleotide retention in actomyosin, such as nucleotide binding and release of products of hydrolysis. To examine the hypothesis of the regulatory role of loop 1 and to determine the mechanism of regulation, multiple chimera studies were performed. The residues in loop 1 were mutated, entire loop 1 was deleted or replaced with loop 1 from other myosins. Obtained results are controversial. In D. discoideum myosin with loop 1 replaced with the one from Acanthamoeba or rabbit myosin, the rate of ADP release from actomyosin correlated with the same parameter of the source myosin (4). No such correlation was observed with smooth myosin (5), it was concluded that the rate of ADP release correlates with the length of loop 1. It was also found that the electrostatics of loop 1 plays a role in the regulation of ADP release from actomyosin (5, 6). Rates of ADP release from actomyosin with β isoforms of the pig and rat cardiac myosin differ about three times, regardless of essentially the same sequence of loop 1 (7). The study concluded that the rate of ADP release from actomyosin is regulated not by loop 1 but by the other parts of the myosin backbone.

**Figure 1.**
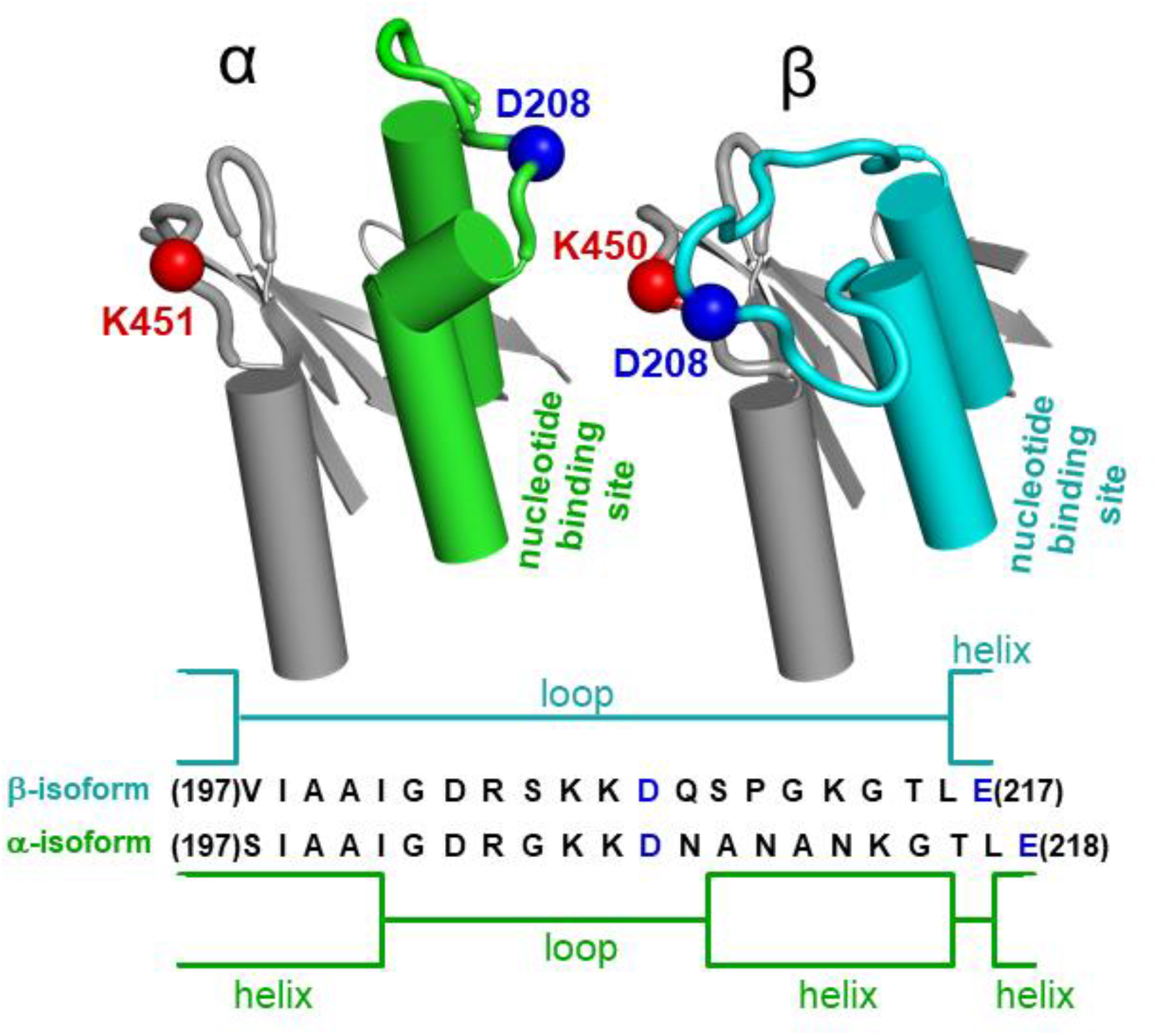
Loop 1 (left, green, α-myosin, and right, cyan, β-myosin) and the 7-stranded β-sheet (gray) are coupled in β myosin and uncoupled in α myosin. Permanent salt bridge D208:K450 electrostatically couples loop 1 and backbone of myosin head. Loop1 is structurally different in the α and β myosin due to the sequence difference ^209^NANAN^213^ (α isoform) and ^209^QSPG^212^ (β isoform), resulting in formation of a short helix in loop 1 of the α isoform. Loop1 in the α myosin is shorter and does not have the salt bridge, which restricts loop 1 flexibility in β myosin. Red and blue spheres are the positive and negative charged residues.

Previously, we run molecular dynamics simulations and analyzed structural dynamics of two isoforms of human cardiac myosin, α and β (8). The sequence homology of the myosin head of these isoforms is 93%, but the sequences of loop 1 are different. The modeling and computer simulations showed that loop 1 in the β isoform forms a salt bridge D208:K450, which restricts loop 1 motion (Figure S1). The salt bridge is absent in the α isoform. The rate of ADP release from actomyosin with the α and β isoforms differ significantly, ADP dissociates from actomyosin with the α isoform faster, than from actomyosin with the β isoform (9). We hypothesize that the salt bridge D208:K450, observed in the β isoform, restricts loop 1 motion and this isoform-specific difference of loop 1 flexibility plays a role in the retention of ADP at the active site. We expressed the WT β isoform of the human cardiac myosin and the D208Q:K450L mutant to remove electrostatic interaction between loop 1 and myosin backbone. We studied the rate of ATP-induced actomyosin dissociation and the rate of ADP release from actomyosin in the WT myosin and mutant. We found that the rates of ADP release from the WT and mutant actomyosin are statistically similar and conclude that the flexibility of loop 1 does not contribute to the regulation of ADP release from actomyosin.

## Materials and Methods

### Ethical approval

Actin was produced from rabbit skeletal tissue. All experimental protocols were approved by the Institutional Animal Care and Use Committee of UNC Charlotte and all experiments were performed in accordance with relevant guidelines and regulations.

### Reagents

*N*-(1-pyrene)iodoacetamide (pyrene) was from Life Technologies Corporation (Grand Island, NY), phalloidin, ATP, and ADP were from Sigma-Aldrich (Milwaukee, WI). All other chemicals were from Fisher Scientific (Waltham, MA) and VWR (Radnor, PA).

### Protein preparation

The β isoform construct of the human cardiac myosin motor domain contains 1-843 residues and a FLAG affinity tag at the C-terminus. Adenoviruses encoded with the wild type and D208Q:K450L myosin mutant were purchased from Vector Biolabs (Malvern, PA), amplified using HEK293 cells (ATCC CRL-1573), and purified using CsCl gradient centrifugation. Recombinant human cardiac myosin was expressed in C_2_C_12_ (ATCC CRL-1722) mouse myoblast cells. C_2_C_12_ cells were grown to a 95% confluence on 15 cm diameter plates and infected with the optimum dosage of virus determined by a viral-titration assay. Cells were allowed to differentiate post-infection and collected seven days post-infection to extract and purify myosin. Collected cells were washed and lysed in the presence of a millimolar concentration of ATP. The cell lysate was incubated with anti-FLAG magnetic beads (Sigma-Aldrich, Milwaukee, WI). Beads were washed and myosin was eluted from the beads by 3x FLAG peptide (ApexBio, Houston, TX). Myosin purity was assessed by coomassie-stained SDS-polyacrylamide gels (Figure S2) and protein concentration was determined by measuring the absorbance at 280 nm using extinction coefficient ε_280nm_ = 93,170 M^−1^cm^−1^, calculated using ProtParam tool of ExPASy web server.

Actin was prepared from rabbit leg and back muscles (10-12). F-actin was labeled with pyrene iodoacetamide (Life Technologies Corporation, Grand Island, NY) with the molar ratio 6:1, label:actin. After labeling, actin was cleaned from the excess of label, re-polymerized, stabilized with phalloidin at the molar ratio of 1:1, and dialyzed for two days at T=4°C against the experimental buffer. Concentration of unlabeled G-actin was determined spectrophotometrically assuming the extinction coefficient ε_290nm_ = 0.63 (mg/ml)^−1^cm^−1^ (13). Concentration of labeled G-actin and labeling efficiency were determined spectroscopically using the following expressions: [G-actin]=(A_290nm_–(A_344nm_·0.127))/26,600 M^−1^ and [pyrene]=A_344nm_/22,000 M^−1^ (14). Actin labeling efficiency was 60% - 80%. The experimental buffer contained 20 mM MOPS (3-[N-morpholino]propanesulfonic acid) pH 7.3, 50 mM KCl, 3mM MgCl_2_. All reported concentrations are final concentrations.

### Acquisition of fluorescent transients

In the ATP-induced actomyosin dissociation experiment, 0.25 µM - 1 µM actomyosin was rapidly mixed with ATP solution of variable concentrations. In the ADP inhibition of the ATP-induced actomyosin dissociation experiment, 0.25 µM - 1 µM actomyosin was rapidly mixed with the premixed ATP and ADP solution. The concentration of ATP in solution was 0.9 mM and the concentration of ADP varied from 10 µM to 200 µM. Transient fluorescence of pyrene-labeled actin was measured with a Bio-Logic SFM-300 stopped flow transient fluorimeter (Bio-Logic Science Instruments SAS, Claix, France), equipped with an FC-15 cuvette. The pyrene fluorescence was excited at 365 nm and detected using a 420 nm cutoff filter. Multiple transients were acquired and averaged to improve the signal-to-noise ratio. 8000 points were acquired in each experiment. All experiments were performed at T=20° C.

### Analysis of fluorescence transients

Obtained transients were fitted with the numerical solution of the system of differential equations, corresponding to the reaction scheme of ATP-induced actomyosin dissociation (Figure 2, Eq.S1) and ADP release from actomyosin (Figure 3, Eq.S2). In both cases, the traces obtained at different nucleotide concentrations for the same protein preparation were fitted simultaneously. Rate constants were obtained in experiments with human cardiac myosin constructs from at least three independent preparations. The transients of the ATP induced actomyosin dissociation were also fitted with the one-exponential function S(t) = S_o_+A·exp(-k_obs_·(t-t_0_)), where S(t) is the observed signal at the time t, A is the signal amplitude, t_0_ is the time before the flow stops, and k_obs_ is the observed rate constant. Transients, obtained for the same actomyosin preparation at different concentrations of the nucleotide were fitted together, assuming the known value of t_0_, measured in a separate experiment, and the constant value of S_0_, which depends on the concentration and labeling efficiency of pyrene-labeled actin in the actomyosin preparation. The dependence of the observed rates k_obs_ on the ATP concentration was fitted by a hyperbola, k_obs_ = V_max_·[ATP]/(K_app_+[ATP]), allowing the determination of the maximum rate, V_max_ (the horizontal asymptote). The rate constant k_+2T_ is the V_max,_ and the equilibrium constant of the collision complex formation K_1T_ is 1/K_app_. To determine the bimolecular rate (K_1T_k_+2T_), the dependence of the observed rates on the ATP concentration was fitted by a straight line at small concentrations of ATP. Transients of the ADP inhibition experiment can be fitted only with a two-exponential function, where the rate constant of the fast process presumably reflects the ATP-induced actomyosin dissociation, and the slower process is governed by ADP release from actomyosin. The fit to the two-exponential function confirms that ADP is not in fast equilibrium with actomyosin, and the reaction (Figure 3, Eq. S2) can be treated as a parallel reaction. Therefore, obtained transients were fitted with the numerical solution of the differential equations, corresponding to the reaction scheme 2, assuming no fast equilibrium of actomyosin and ADP. All exponential data fits were performed with Origin 8 (OriginLab Corp, Northampton MA). The statistical significance of results was tested with ANOVA integrated into Origin 8 software. A significance level of P < 0.05 was used for all analyses. Differential equations were solved numerically using Wolfram Mathematica built-in symbol NDSolve. The solution was fitted to the experimental data using the built-in symbol NMinimize, which searches for a global minimum. Wolfram Mathematica scripts can be found in the Supporting Material.

**Figure 2.**
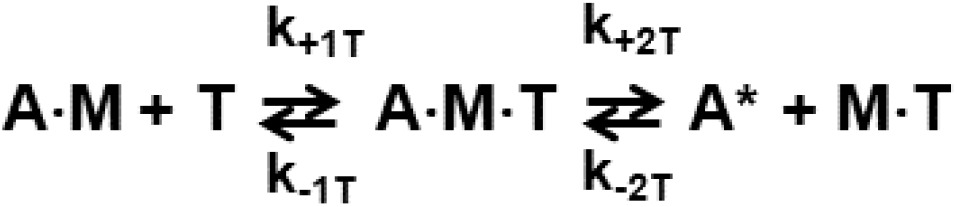
Scheme of the ATP-induced actomyosin dissociation. A = pyrene-labeled actin, M = myosin, T = ATP. A* = actin with unquenched pyrene fluorescence.

**Figure 3.**
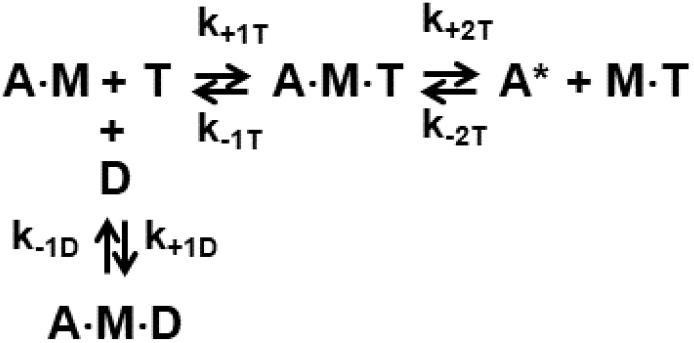
Scheme of the parallel reaction of ATP or ADP binding to actomyosin. A = pyrene-labeled actin, M = myosin, T = ATP. A* = actin with unquenched pyrene fluorescence.

**Figure 4.**
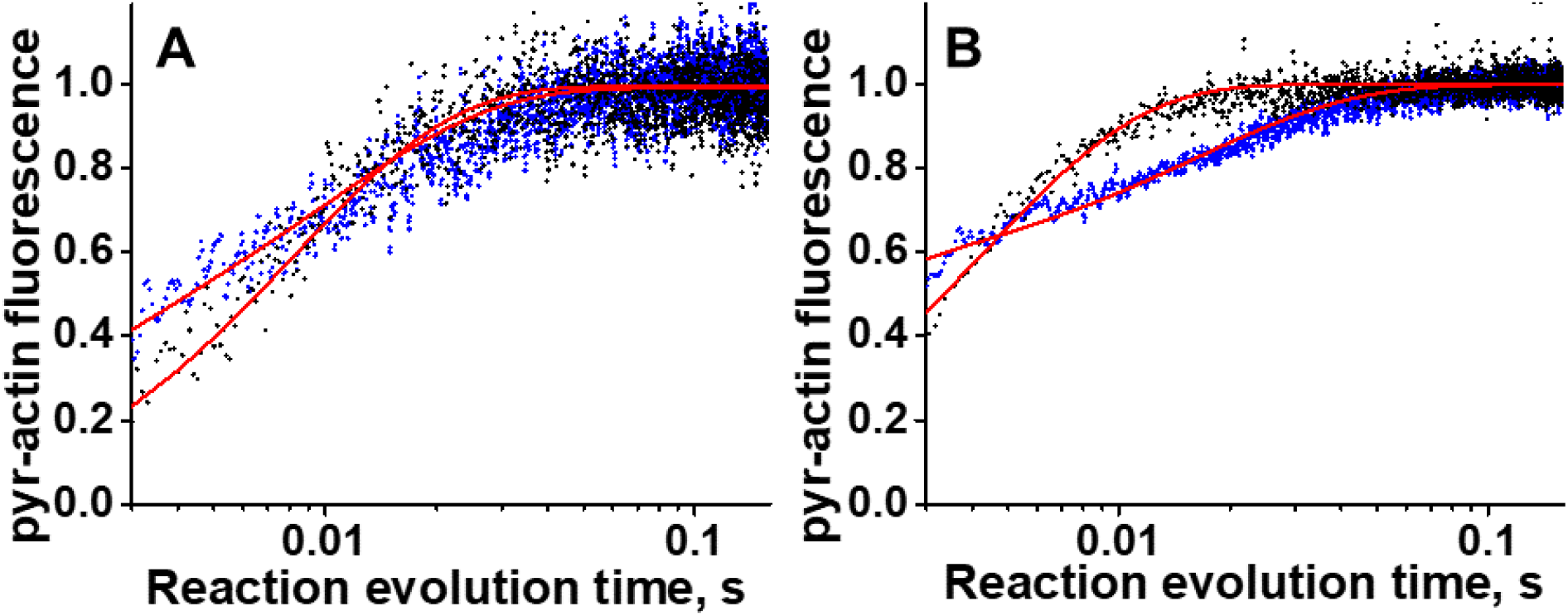
Typical transients obtained in the study. Pyrene labeled actomyosin is rapidly mixed with ATP (60 μM, black dots), or the mixture of ATP and ADP ([ATP]=900 μM, [ADP]=100 μM, blue dots). Normalized pyrene actin fluorescence reflects actomyosin dissociation. The transients are fitted to the numerical solution of differential equations (Eq. S1, S2), red traces. All transients obtained for the same myosin preparation are fitted globally with the same set of kinetic constants. Different signal to noise ratio reflects different labeling efficiency of actin. A, WT myosin, B, D208Q:K450L myosin mutant. The rate of actomyosin dissociation is faster in the mutant and larger portion of transient is hidden in the dead time of the spectrofluorometer.

## Results

### Mutation D208Q:K450L decreases the lifetime of ATP-actomyosin collision complex and increases the rate of actomyosin dissociation

ATP binding and actomyosin dissociation were monitored using fluorescence of pyrene labeled actin. Pyrene fluorescence is quenched when actin strongly binds myosin and increases upon actomyosin dissociation. In the case of the WT myosin the maximum rate of actomyosin dissociation was achieved at 450 μM - 900 μM ATP. In the case of the mutant the rate of actomyosin dissociation is higher and at ATP concentration of 150 μM and higher the transients were flat, indicating that the actomyosin dissociation occurs during the dead time of the stopped flow fluorometer. Therefore, for the mutant we run the reaction at smaller concentrations of ATP (15 μM – 100 μM). In the assumption of the rapid equilibrium of the ATP-actomyosin collision complex formation (step 1 in Figure 2) we fit obtained transients with single-exponential function to determine rection rates at different concentrations of ATP, and the rates were fitted with a hyperbola to determine V_max_ and K_app_ (Figure S3). The rate constant K_1T_k_+2T_ was determined from the slope of the linear fit of the dependence of measured reaction rates on ATP concentration (Figure S3). We also fitted obtained transients with the numerical solution of the system of differential equations corresponding to the reaction scheme of ATP-induced actomyosin dissociation (Figure 2). For the WT actomyosin, both fits produced similar values for the rate constant k_+2T_, 491.5 ± 74.1 s^−1^ for the fit with a hyperbola, and 406.2 ± 14.1 s^−1^ for the fit with differential equations. The values for K_1T_k_+2T_ were about two times different in the fits (2.1 ± 0.2 μM^−1^s^−1^ and 4.6 ± 1.0 μM^−1^s^−1^ for the fit with a hyperbola and with differential equations respectively), the fit with differential equations suggests tighter binding of ATP and actomyosin in the collision complex than the fit of transients with exponential function. For the mutant both fits produced similar results, k_+2T_ = 866.4 ± 317.3 s^−1^ and 961.0 ± 97.1 s^−1^, K_app_ = 200.9 ± 94.0 μM and 238.9 ± 106.5 μM, K_1T_k_+2T_ = 3.6 ± 0.1 μM^−1^s^−1^ and 4.3 ± 2.0 μM^−1^s^−1^ for the fit with a hyperbola and with differential equations respectively. Data for the WT actomyosin were taken from our previous study (15). Obtained rate constants k_+1T_ and k_-2T_ are statistically similar for the WT and mutant (k_+1T_ = 7.4 ± 1.4 µM^−1^s^−1^ (WT) and 9.9 ± 1.0 µM^−1^s^−1^ (mutant), k_-2T_ = 3.6 ± 2.5 s^−1^ (WT) and 0.9 ± 0.9 s^−1^ (mutant), (Table 1)). Due to discrepancy of the fits for the WT actomyosin and the mutant, and due to the high standard deviation of the rate for the mutant, it is difficult to conclude if the rate constant k_-1T_ is different in the WT and mutant actomyosin. The rate constants k_+2T_ are statistically different for the WT and mutant (k_+2T_ = 406.2 ± 14.1 s^−1^ (WT) and 961.0 ± 97.1 s^−1^ (mutant), (Table 1)). Our data show that D208Q:K450L mutation leads to the increased rate of ATP-induced actomyosin dissociation.

**Table 1.**
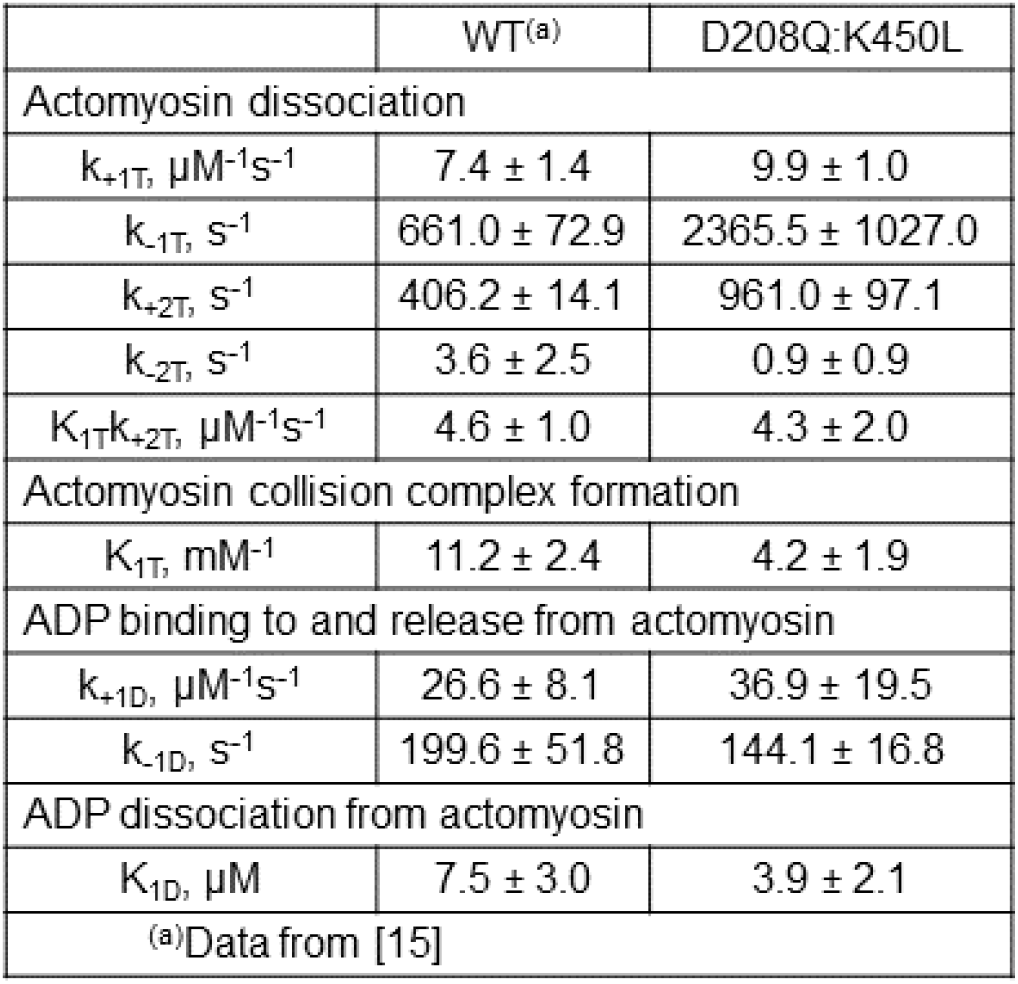
Actomyosin kinetic rate constants, obtained in the fit of the transients with the numerical solution of differential equations Eqs. S1 and S2, mean ± SD. Data are averages of three (WT) and five (mutant) independent protein preparations

### Mutation D208Q:K450L does not affect the rate of ADP release from actomyosin

ADP release from actomyosin was monitored via the change of pyrene fluorescence, assuming that ADP release is a slower process than the ATP-induced actomyosin dissociation. In the experiment, actomyosin was rapidly mixed with the premixed ATP and ADP. We kept concentration of ATP constant and varied ADP concentration. Previously we found that ADP has a high affinity to actomyosin with human cardiac myosin (15), and obtained transients can be fitted with a double exponential function, confirming that ADP is not in fast equilibrium with actomyosin. We fitted obtained transients to the solution of the differential equations, corresponding to the reaction scheme of parallel reaction, without assumption of the fast equilibrium of actomyosin and ADP upon rapid mixing. Traces, obtained at different concentrations of ADP were fitted simultaneously to determine the reaction rate constants (Table 1). We found that for the WT and mutant, the rate constants k_+D_ and k_-D_ are statistically similar (k_+D_ = 26.6 ± 8.1 µM^−1^s^−1^ (WT) and 36.9 ± 19.5 µM^−1^s^−1^ (mutant), k_-D_ = 199.6 ± 51.8 s^−1^ (WT) and 144.1 ± 16.8 s^−1^ (mutant), (Table 1)). Our data show that D208Q:K450L mutation does not affect the rate of ADP dissociation from actomyosin.

## Discussion

ATP binding to actomyosin is a two-step process, formation of the weak collision complex, and then the tight binding, which results in actomyosin dissociation. Two loops at the active site play a role in ATP binding to actomyosin, P-loop and loop switch 1. According to the current view, ATP first binds P-loop and then loop switch 1 closes to stabilize and coordinate MgATP in the active site. This motion of loop switch 1 results in actomyosin dissociation. Another loop of the active site, loop switch 2, subsequently closes and interacts with loop switch 1 to ensure successful ATP hydrolysis. After ATP hydrolysis, abstracted phosphate is released via the “trapdoor” by destabilization and opening of loop switch 1 (16). The fluorescence of pyrene actin reflects a transition from the strong to weak actomyosin binding, therefore the fluorescence change due to actomyosin dissociation can be interpreted as an indicator of the transition of loop switch 1 from the open to closed state. According to our data, mutation D208Q:K450L does not affect the rate of ATP binding to the active site (k_+1T_). The rate of actomyosin dissociation (k_+2T_) is two times faster in the mutant. The rates of ADP binding and dissociation (k_+D_ and k_-D_) are similar in the WT and mutant actomyosin.

Obtained rate constants of nucleotide binding to actomyosin k_+1T_ and k_+D_ are lower than the theoretical rate of the diffusion-limited nucleotide binding (255 µM^−1^s^−1^, (15)). Both MgATP and MgADP complexes possess electric charge ((MgATP)^2-^ and (MgADP)^-^, (17)), therefore, we conclude that electrostatic repulsion governs the rate of nucleotide binding, and the difference of the reaction rates k_+1T_ and k_+D_ reflects the charge difference of MgATP and MgADP. In the WT actomyosin, the rate of ATP dissociation from the collision complex k_-1T_ is higher than the rate of ADP dissociation k_-D_. Assuming that the formation of the collision complex reflects only nucleotide binding to P-loop, one can suggest that the faster rate k_-1T_ is the consequence of the charge difference of these nucleotides.

Our study shows that the rate k_+2T_ is higher in the mutant, and the rate k_-D_ is not affected by the mutation. According to the crystal structures of myosin with ADP and with ATP analogs, loop switch 1 is closed when nucleotide is bound to the active site. When ATP analog is bound to the active site, P-loop coordinates γ and β phosphates and Mg ion, and loop switch 1 coordinates Mg ion and γ phosphate. When ADP is bound, all γ phosphate coordinations are lost, and loop switch 1 coordinates only Mg ion when P-loop coordinates Mg ion and β phosphate of ADP. Faster rate k_+2T_ could indicate the increased dynamics of relative positions of loop switch 1 and P-loop in the mutant. Such an increased flexibility decreases time of loop switch 1 closure and coordination of MgATP in the active site for successful hydrolysis (faster rate k_+2T_). The rate k_-D_ is similar in the WT and mutant, and this similarity indicates that the coordination of ADP by P-loop governs kinetics of ADP and actomyosin interaction. There is no direct interaction between ADP and loop switch 1, and therefore the increased flexibility of loop switch 1 does not affect the rate of ADP dissociation. According to the “trapdoor” mechanism, loop switch 1 opens upon ATP hydrolysis, therefore, the interaction of loop switch 1 with Mg ion is weak, it does not stabilize loop switch 1 in the closed state and does not affect the retention of ADP in the active site. Thus, our data show that the electrostatic interaction of loop 1 with the backbone of myosin head affects flexibility of P-loop and loop switch 1 at the active site. Relative flexibility of loop switch 1 and P-loop at active site affects ATP binding and the rate of actomyosin dissociation, but it does not affect the rate of ADP dissociation from human cardiac actomyosin.

## Acknowledgements

Funding: This work was supported by the National Institutes of Health [grant number HL132315].

## References

1. Murphy CT, Spudich JA. Variable surface loops and myosin activity: accessories to a motor. J Muscle Res Cell Motil. 2000;21(2):139–51. Epub 2000/08/29. doi: 10.1023/a:1005610007209. PubMed PMID: 10961838.

2. Furch M, Geeves MA, Manstein DJ. Modulation of actin affinity and actomyosin adenosine triphosphatase by charge changes in the myosin motor domain. Biochemistry. 1998;37(18):6317–26. Epub 1998/06/13. doi: 10.1021/bi972851y. PubMed PMID: 9572846.

3. Spudich JA. How molecular motors work. Nature. 1994;372(6506):515–8. Epub 1994/12/08. doi: 10.1038/372515a0. PubMed PMID: 7990922.

4. Murphy CT, Spudich JA. Dictyostelium myosin 25-50K loop substitutions specifically affect ADP release rates. Biochemistry. 1998;37(19):6738–44. Epub 1998/06/06. doi: 10.1021/bi972903j. PubMed PMID: 9578557.

5. Sweeney HL, Rosenfeld SS, Brown F, Faust L, Smith J, Xing J, Stein LA, Sellers JR. Kinetic tuning of myosin via a flexible loop adjacent to the nucleotide binding pocket. J Biol Chem. 1998;273(11):6262–70. Epub 1998/04/16. doi: 10.1074/jbc.273.11.6262. PubMed PMID: 9497352.

6. Clark R, Ansari MA, Dash S, Geeves MA, Coluccio LM. Loop 1 of transducer region in mammalian class I myosin, Myo1b, modulates actin affinity, ATPase activity, and nucleotide access. J Biol Chem. 2005;280(35):30935–42. Epub 2005/06/28. doi: 10.1074/jbc.M504698200. PubMed PMID: 15980431.

7. Pereira JS, Pavlov D, Nili M, Greaser M, Homsher E, Moss RL. Kinetic differences in cardiac myosins with identical loop 1 sequences. J Biol Chem. 2001;276(6):4409–15. Epub 2000/11/15. doi: 10.1074/jbc.M006441200. PubMed PMID: 11076938.

8. Gargey A, Ge J, Tkachev YV, Nesmelov YE. Electrostatic interactions in the force-generating region of the human cardiac myosin modulate ADP dissociation from actomyosin. Biochem Biophys Res Commun. 2019;509(4):978–82. Epub 2019/01/19. doi: 10.1016/j.bbrc.2019.01.045. PubMed PMID: 30654937; PMCID: PMC6348005.

9. Deacon JC, Bloemink MJ, Rezavandi H, Geeves MA, Leinwand LA. Identification of functional differences between recombinant human alpha and beta cardiac myosin motors. Cell Mol Life Sci. 2012;69(13):2261–77. Epub 2012/02/22. doi: 10.1007/s00018-012-0927-3. PubMed PMID: 22349210; PMCID: PMC3375423.

10. Margossian SS, Lowey S. Preparation of myosin and its subfragments from rabbit skeletal muscle. Methods Enzymol. 1982;85 Pt B:55–71. PubMed PMID: 6214692.

11. Strzelecka-Golaszewska H, Prochniewicz E, Nowak E, Zmorzynski S, Drabikowski W. Chicken-gizzard actin: polymerization and stability. Eur J Biochem. 1980;104(1):41–52. PubMed PMID: 6445264.

12. Waller GS, Ouyang G, Swafford J, Vibert P, Lowey S. A minimal motor domain from chicken skeletal muscle myosin. J Biol Chem. 1995;270(25):15348–52. PubMed PMID: Medline:7797523.

13. Houk TW, Jr., Ue K. The measurement of actin concentration in solution: a comparison of methods. Anal Biochem. 1974;62(1):66–74. Epub 1974/11/01. doi: 0003-2697(74)90367-4 [pii]. PubMed PMID: 4473917.

14. Takagi Y, Yang Y, Fujiwara I, Jacobs D, Cheney RE, Sellers JR, Kovacs M. Human myosin Vc is a low duty ratio, nonprocessive molecular motor. J Biol Chem. 2008;283(13):8527–37. doi: 10.1074/jbc.M709150200. PubMed PMID: 18201966.

15. Gargey A, Iragavarapu SB, Grdzelishvili AV, Nesmelov YE. Electrostatic interactions in the SH1-SH2 helix of human cardiac myosin modulate the time of strong actomyosin binding. J Muscle Res Cell Motil. 2020. Epub 2020/09/16. doi: 10.1007/s10974-020-09588-1. PubMed PMID: 32929610; PMCID: PMC7956043.

16. Reubold TF, Eschenburg S, Becker A, Kull FJ, Manstein DJ. A structural model for actin-induced nucleotide release in myosin. Nat Struct Biol. 2003;10(10):826–30. Epub 2003/09/23. doi: 10.1038/nsb987. PubMed PMID: 14502270.

17. Cowan JA. Metallobiochemistry of Magnesium. Coordination Complexes with Biological Substrates: Site Specificity, Kinetics and Thermodynamics of Binding, and Implications for Activity. Inorg Chem. 1991;2740-2747:2740–7.

