## Supplementary Information for "Loop 1 of human cardiac myosin regulates the rate of ATP induced actomyosin dissociation but not the rate of ADP release"

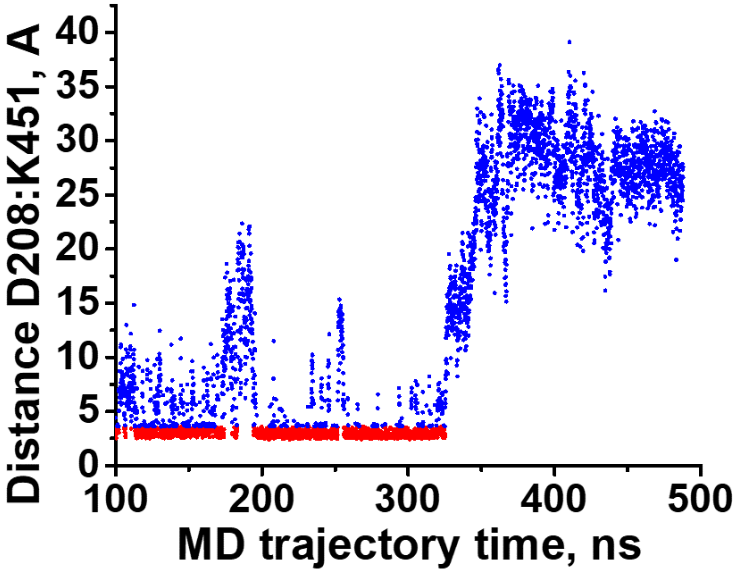


**Figure S 1.** Salt bridge D208:K451 in the loop1 region of the α isoform. Red dots indicate a salt bridge (the distance between carboxylic oxygen (negative charge, H acceptor) and amino nitrogen (positive charge, H donor) is less than 3.6 A), blue dots show the absence of the bridge. Since the template for a homology model of the α isoform was the β isoform, the α isoform had the bridge in the beginning of the molecular dynamics simulation. After approx. 325 ns of the simulation, the helix A210:T216 forms and the bridge breaks permanently.

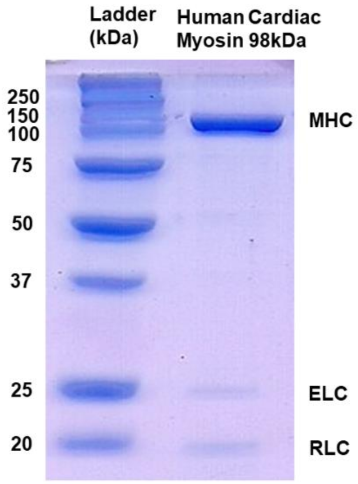


**Figure S 2**. SDS-PAGE of the purified recombinant myosin head. 98 kDa human cardiac S1 co-purifies with murine ELC and RLC

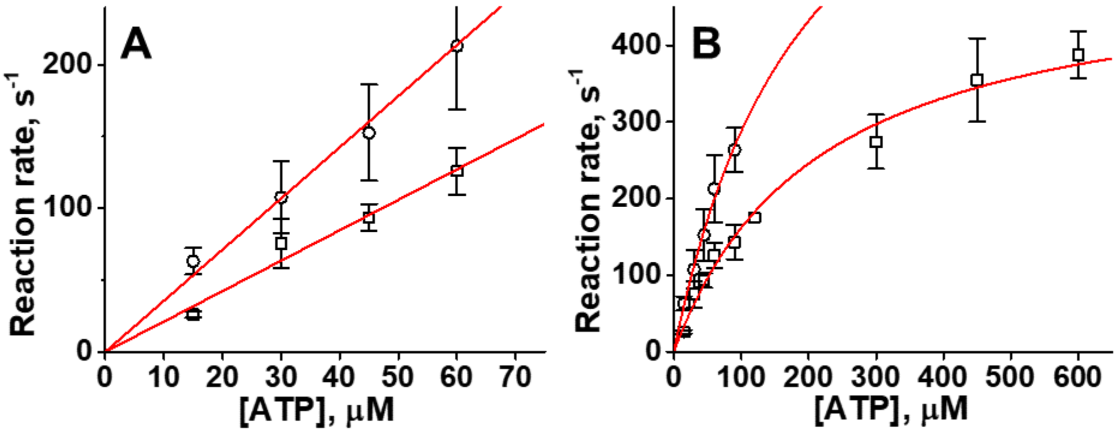


**Figure S 3.** ATP-induced actomyosin dissociation. Squares, WT, data from [], N = 3. Circles, mutant, N = 5. A. Observed reaction rates at low [ATP], fitted with a straight line, the second-order reaction rate constant is determined from the slope of the line. $K_{1T}k_{+2T}$ = 2.12 ± 0.21 µM-1s−1 and 3.56 ± 0.13 µM−1s−1 for WT and mutant, respectively. B. Reaction rates fitted with a hyperbola. For the mutant, the reaction rate is too fast for accurate measure above [ATP] = 100 µM. Rates k+2T = 491.5 ± 74.1 s−1 and 866.4 ± 317.3 s−1 for WT and mutant, respectively. Equilibrium dissociation constant $K_{app}$ = 214.2 ± 24.9 µM and 200.9 ± 94.0 µM. Data points are mean ± SD. N is the number of biological replicates.

Differential equations were solved numerically using Wolfram Mathematica built-in symbol NDSolve. The solution was fitted to the experimental data using the built-in symbol NMinimize, which searches for a global minimum. Maximum number of iterations was usually set to 100. If minimization was not completed, maximum 200 iterations were used. We first fitted transients from the experiment with [ADP] = 0 (Scheme 1) to determine reaction rate constants $k_{+1T}$, $k_{-1T}$, $k_{+2T}$, $k_{-2T}$. Transients obtained from the same myosin construct preparation were fitted globally. We used obtained reaction rate constants in the fit of transients, obtained in experiments with [ADP] ≠ 0. All transients obtained in reactions using myosin from the same preparation were fitted globally for all used ADP concentrations. Reaction rate constants determined from the fits for different preparations of the same myosin construct were averaged and reported as mean ± standard deviation.

1. Differential equations corresponding to the reaction shown in Scheme 1.

Eq. S1

$$\frac{dAM}{dt} = -k_{+1T}AM \cdot T + k_{-1T}AMT$$

$$\frac{dAMT}{dt} = k_{+1T}AM \cdot T - k_{-1T}AMT - k_{+2T}AMT + k_{-2T}A \cdot MT$$

$$\frac{dMT}{dt} = k_{+2T}AMT - k_{-2T}A \cdot MT$$

$$\frac{dA}{dt} = k_{+2T}AMT - k_{-2T}A \cdot MT$$

2. Wolfram Mathematica script to solve the system of differential equations Eq. S1 numerically and fit obtained transient to the numerical solution. Input: data (ASCII file with the experimental data). Output: rate constants $k_{+1T}$, $k_{-1T}$, $k_{+2T}$, $k_{-2T}$.

```
sse[kp1T_?NumberQ,km1T_?NumberQ,kp2T_?NumberQ,km2T_?NumberQ]:=Block[{sol},sol=NDSolve[{
am'[x]==-kp1T*am[x]*T+km1T*amt[x],
amt'[x]==kp1T*am[x]*T-(km1T+kp2T)*amt[x]+km2T*a[x]*mt[x],
a'[x]==kp2T*amt[x]-km2T*a[x]*mt[x],
mt'[x]==kp2T*amt[x]-km2T*a[x]*mt[x],
am[0]==0.5, amt[0]==0,a[0]==0,mt[0]==0},{a},{x,0,0.8}][[1]];Plus@@Apply[(a[#1]-#2)^2&,data,{1}]/.sol]
NMinimize[{sse[kp1T,km1T,kp2T,km2T],0.5<=kp1T<=80,100<=km1T<=3500,50<=kp2T<=1600,0.001<=km2T<=50},{
{kp1T,1,15},{km1T,300,500},{kp2T,500,600},{km2T,1,2}}]
```

3. Differential equations corresponding to the reaction shown in Scheme 2.

$$\frac{dAM}{dt} = -k_{+1T}AM \cdot T + k_{-1T}AMT - k_{+1D}AM \cdot D + k_{-1D}AMD \qquad \text{Eq. S2}$$
$$\frac{dAMT}{dt} = k_{+1T}AM \cdot T - (k_{-1T} + k_{+2T})AMT + k_{-2T}AM \cdot T$$
$$\frac{dA}{dt} = k_{+2T}AMT - k_{-2T}A \cdot MT$$
$$\frac{dMT}{dt} = k_{+2T}AMT - k_{-2T}A \cdot MT$$
$$\frac{dAMD}{dt} = k_{+1D}AM \cdot D - k_{-1D}AMD$$
$$\frac{dD}{dt} = -k_{+1D}AM \cdot D + k_{-1D}AMD$$

4. Wolfram Mathematica script to solve the system of differential equations Eq. S2 numerically and fit the obtained transient to the numerical solution. Input: data (ASCII file with the experimental data), rate constants $k_{+1T}$, $k_{-1T}$, $k_{+2T}$, $k_{-2T}$, determined in the previous fit. Output: rate constants $k_{+1D}$, $k_{-1D}$.

```
sse[kp1D_?NumberQ,km1D_?NumberQ]:=Block[{sol},sol=NDSolve[{
    am'[x]==-kp1T*am[x]*T+km1T*amt[x]-kp1D*am[x]*d[x]+km1D*amd[x],
    amt'[x]==kp1T*am[x]*T-(km1T+kp2T)*amt[x]+km2T*a[x]*mt[x],
    a'[x]==kp2T*amt[x]-km2T*a[x]*mt[x],
    mt'[x]==kp2T*amt[x]-km2T*a[x]*mt[x],
    amd'[x]==kp1D*am[x]*d[x]-km1D*amd[x],
    d'[x]==-kp1D*am[x]*d[x]+km1D*amd[x],
    am[0]==0.5, amt[0]==0,a[0]==0,mt[0]==0,d[0]==ADP,amd[0]==0},{a},{x,0,0.8},][[1]];Plus@@Apply[(a[#1]-
#2)^2&,data,{1}]/.sol]
NMinimize[{sse[kp1D,km1D],1<=kp1D<=200,10<=km1D<=800},{{kp1D,1,10},{km1D,100,200}}]
```
